# Dietary fat promotes antibiotic-induced *Clostridioides difficile* mortality in mice

**DOI:** 10.1101/828939

**Authors:** Keith Z. Hazleton, Casey G. Martin, David J Orlicky, Kathleen L. Arnolds, Nichole M. Nusbacher, Nancy Moreno-Huizar, Michael Armstrong, Nichole Reisdorph, Catherine A. Lozupone

**Author notes:** Department of Pediatrics, Division of Gastroenterology, Hepatology and Nutrition, University of Arizona, Tucson, AZ 85719.

## Abstract

*Clostridioides difficile* infection (CDI), is the leading cause of hospital-acquired diarrhea and emerging evidence has linked dietary components with CDI pathogenesis, suggesting that dietary modulation may be an effective strategy for prevention. Here, we show that mice fed a high-fat/low-fiber “Western type” diet (WD) had dramatically increased mortality in a murine model of antibiotic-induced CDI compared to a low-fat/low-fiber (LF/LF) diet and standard mouse chow controls. We found that the WD had a pro- *C. difficile* bile acid composition that was driven in part by higher levels of primary bile acids that are produced to digest fat, and a lower level of secondary bile acids that are produced by the gut microbiome. This lack of secondary bile acids was associated with a greater disturbance to the gut microbiome with antibiotics in both the WD and LF/LF diet compared to mouse chow. Mice fed the WD also had the highest level of toxin TcdA just prior to the onset of mortality, but not of TcdB or increased inflammation. These findings indicate that dietary intervention to decrease fat may complement previously proposed dietary intervention strategies to prevent CDI in high-risk individuals.

**One Sentence Summary:** A high-fat/low-fiber Western type diet promoted mortality in a mouse model of antibiotic-induced *C. difficile* infection compared to a low-fat/low-fiber diet and chow diet, suggesting that lower dietary fat may be an effective strategy for preventing *C. difficile* pathology.

## Introduction

*Clostridioides difficile* infection (CDI) is an important cause of morbidity and mortality, with 500,000 cases every year causing 30,000 deaths per year in the US alone^1^. Alarmingly, there has been a steady increase in the number of new infections in spite of prevention efforts in hospitals that have focused largely on increased sanitation and antibiotic stewardship^2^. *C. difficile* induced pathology has been linked to the production of two different toxins, TcdA and TcdB, which can directly induce intestinal damage and inflammation ^3^.

A complex gut microbiome is protective against CDI^4^. Illnesses associated with reduced gut microbiome diversity, such as inflammatory bowel disease^5^ increase risk of CDI, as does broad spectrum antibiotic usage, such as clindamycin, beta-lactams, and fluoroquinolones ^6, 7^. Antibiotics have been shown to predispose mice to CDI via modified metabolic activity of the altered gut microbiome ^8^. Individuals with recurrent CDI (rCDI) typically have microbiomes with greatly reduced complexity and altered composition ^9–11^. The gut microbiome provides protection from CDI in part through metabolism of primary bile acids, which are excreted by the liver into the intestine where they play a central role in fat digestion^12^. The primary bile acids taurocholic acid (TCA) and cholic acid (CA) can promote the germination of *C. difficile* spores. However, a healthy gut microbiome can metabolize TCA and CA into the secondary bile acid deoxycholate (DCA), a derivative that can arrest the growth of vegetative *C. difficile* ^13^. Accordingly, prior studies have shown that secondary bile acid producers such as *Clostridium scindens* can protect against CDI in mice^14^. Short chain fatty acids (SCFA), which are microbial products of fermentation of dietary microbial accessible carbohydrates (MACs; e.g. soluble fibers such as inulin), have also been shown to directly suppress *C. difficile* growth *in vitro* ^15^ and are decreased in individuals with rCDI ^16^.

Recent studies conducted in mouse models of antibiotic-induced CDI have suggested that diet modulation has the potential to be an effective prevention strategy for antibiotic-induced CDI. Diets high in MACs^15^ and low in proline ^4^ reduced *C. difficile* colonization and persistence. Excess dietary zinc reduces the threshold of antibiotics needed to confer susceptibility to CDI and increased cecal inflammation and toxin activity ^17^. High-fat/high-protein diets ^18^ and high-fat induced obesity ^19^ resulted in more severe disease and/or increased mortality. Furthermore, mice fed a protein deficient defined diet, had increased survival, decreased weight loss, and decreased overall disease severity ^20^.

Given that primary bile acids play a central role in fat digestion, increase with diets high in saturated fat ^21^ and are a germination factor for *C. difficile* spores, we became interested in investigating a role for dietary fat in antibiotic-induced CDI pathogenesis. We hypothesized that a high-fat diet coupled with low-fiber in the context of antibiotic treatment would provide a “double hit” for shifting towards a pro-*C. difficile* bile acid pool – with dietary fat increasing excretion of pro- *C. difficile* primary bile acids into the gut and increased antibiotic-induced gut microbiome disturbance decreasing their conversion into protective secondary bile acids. We found that high dietary fat content in the context of a low fiber diet (a high-fat/low-fiber Western Diet; WD) induced high mortality from CDI in an antibiotic-induced *C. difficile* model. This higher mortality was linked with higher levels of *C. difficile* toxin TcdA, but not with higher levels of TcdB or increased intestinal inflammation just prior to the onset of mortality. Bile acid pools were strongly influenced by diet to a pro-*C. difficile* composition, but more work needs to be done to determine the degree to which these differences were driving the higher mortality observed with the WD. Our work suggests that dietary interventions to decrease fat intake may complement previously proposed strategies that target fiber and protein to prevent CDI in high-risk individuals.

## Results

### High dietary fat in the context of low dietary fiber causes increased mortality in murine antibiotic-induced CDI

To understand the effects of dietary fat on CDI, we used an established murine model of antibiotic-induced CDI ^22^. Specifically, conventional 6-week-old old female C57BL/6 mice were fed 1 of 3 diets: 1) conventional mouse chow that is low-fat/high-fiber, 2) a purified “Western” diet (WD) that had ∼2x the content of fat with increased ratio of saturated-to-unsaturated fat compared to chow and only insoluble cellulose as a source of fiber, and 3) a similar purified diet as the WD, but with a lower fat content, similar to chow (low-fat/low-fiber; LF/LF) (Table 1; Table S2). This WD composition represents a typical diet in the United States based on population survey data with 34.5% of calories from fat, with a roughly equivalent contributions of saturated (∼36%), mono-unsaturated fats (41%) and a lower contribution from poly-unsaturated fats (∼21%). In the LF/LF diet, these contributions are reversed (saturated fat ∼19% and poly-unsaturated fat ∼39%). One week after diet switch, mice were treated with a cocktail of antibiotics in their drinking water for 5 days (kanamycin, gentamicin, colistin, metronidazole and vancomycin) followed by an injection of clindamycin and gavage with *C. difficile* VPI 10463 (Fig. 1A). The experiments were carried out for up to 21 days past *C. difficile* gavage, allowing us to assay effects on mortality and relate these to fecal microbiome composition. Experiments were conducted on 2 separate cohorts – 2 cages of five mice per diet, with technical replicates of mice with a total sample size of 20 per diet (Figure 1, Table S2).

**Figure 1:**
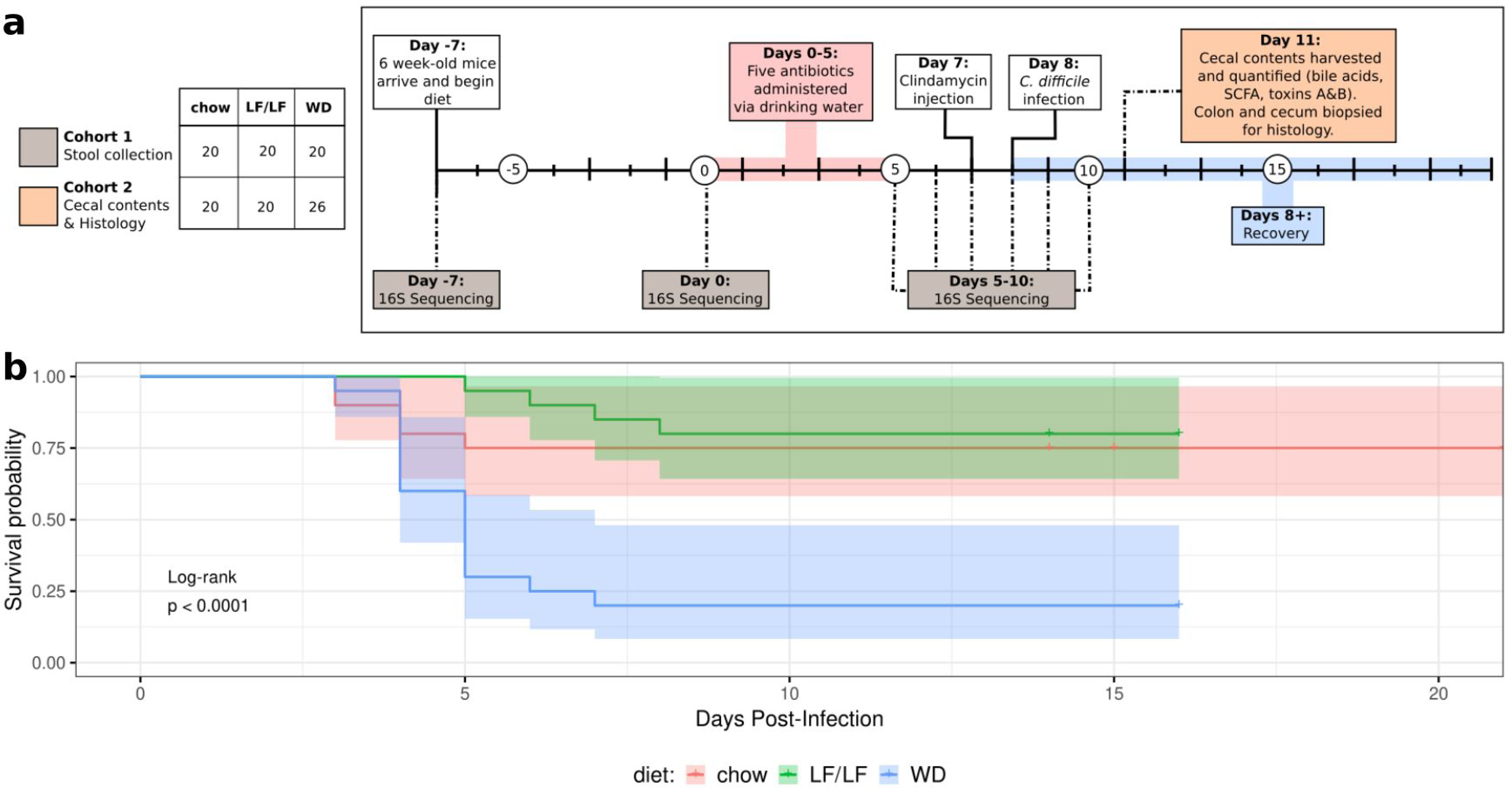
Experimental design of murine model of antibiotic-induced CDI and survival curves. **(A)** *C. difficile* challenge experimental design. The figure legend at the left panel indicates the samples sizes for 2 cohorts; more information on batching and n’s per assay is given in Table S2). Cohort 1 was followed for 13 days post *C. difficile* gavage to monitor survival and gut microbiome composition over time. Cohort 2 was sacrificed at 3 days post *C. difficile* gavage to collect cecal contents for measurement of metabolites and toxin and colon and cecal mucosa for histopathology (some assays were only conducted on a subset of Cohort 2; but all in at least 2 independent experiments; see Table S2 for details). Grey and orange boxes indicate the timepoints at which samples were collected for the respective cohorts. (**B)** Survival curves on the 3 diets. Statistical significance as assessed by log-rank comparison is indicated.

**Table 1.** Diet Composition

Diet Composition
|  | **Chow diet** | **WD** | **LF/LF diet** |
| --- | --- | --- | --- |
| Fat (% kcal)<br>(% *SFA*)<br>(% *MUFA*)<br>(% *PUFA*) | 16<br>(*N/A*) | 34.5<br>(*36.2*)<br>(*41.3*)<br>(*22.5*) | 17.2<br>(*19.5*)<br>(*41.7*)<br>(*38.8*) |
| Carbohydrates (% kcal)<br>(*Sucrose*) | 60<br>(*0*) | 50<br>(*23.4*) | 63.9<br>(*10.6*) |
| Protein (%kcal) | 24 | 15.5 | 18.8 |
| Fiber (g/kg) | 137 | 50 (cellulose) | 50 (cellulose) |

The WD-fed mice showed a marked increase in mortality as compared to both the LF/LF (HR 7.403 p = 0.0041) and chow-fed mice (HR 4.95 p = 0.00208) upon *C. difficile* exposure.

Mortality onset began at Day 4 in the WD and chow-fed mice and then continued to Day 8 in the WD and to Day 6 in the chow-fed mice before stabilizing with the remaining mice appearing to recover. The LF/LF diet-fed mice showed survival levels comparable to the chow-fed mice with a slightly delayed onset of mortality (Fig. 1B). WD-fed diet mice did not show increased weight loss compared to the other diets (Figure S1). Qualitatively, WD-fed mice had more purulent and liquid stools, and poorer grooming than LF/LF and chow-fed mice starting 2 days after infection. Because our WD and LF/LF diet differed in sucrose content, we also tested a fourth diet that was low in fat and fiber, but with sucrose equivalent to the WD (Table S1). Sucrose did not appear to play a role in the increased mortality observed in the WD, as 100% survival was observed in mice fed the low-fat/low-fiber/low-sucrose diet. (n = 10, one cage with 5 mice in two separate experiments).

### WD associated with increased C. difficile toxin TcdA but not TcdB or intestinal inflammation

To further explore the mechanisms of increased mortality in the WD-fed mice by assessing factors that required the collection of host tissues, we conducted a second set of experiments in which mice were sacrificed at day 3 post *C. difficile* gavage (cohort 2; Figure 1). We chose 3 days post *C. difficile* gavage because this was just prior to the observed onset of mortality in the first cohort across all 3 diets (Fig. 1B), and because we felt it was important to compare all mice at a standard time point.

We measured cecal levels of *C. difficile* Toxins A (TcdA) and B (TcdB) by ELISA (see Table S2 for cohort size and batch information). Interestingly, TcdA and not TcdB showed differences with diet consistent with mortality patterns, with TcdA being much higher in the WD compared to the LF/LF diet. The LF/LF diet also had slightly lower levels of TcdA than the chow diet (Figure 2A), which is consistent with a delayed onset of mortality in the LF/LF diet compared to chow (Figure 1). To understand whether intestinal inflammation was related to toxin levels and differed between diets, we evaluated the transverse colon and cecum by histology (Fig. 2B, D). Cecal and transverse colon tissues from mice sacrificed three days post-infection with *C. difficile* were fixed and stained with hematoxylin and eosin and were scored by the Barthel and Dieleman scoring systems respectively by a trained histologist blinded to the treatments and grouping of individuals ^23, 24^. The cecum and distal colon samples showed mild to moderate inflammation, but the histologic damage did not differ across diet groups (Fig. 2B; representative histology Fig. 2D). To control for batch effects, we also assessed differences between diet groups for both toxins and cecal/colon inflammation with linear regression that included batch in the model (Table S2; Figure S2). These results were similar but showed significantly increased inflammation in chow-fed versus WD-fed mice (Figure S2). Cecal levels of TcdB and not TcdA strongly correlated with cecal inflammation (Figure 2; Table S3). Taken together, our results suggest that although TcdB levels are associated with higher intestinal inflammation in these mice just prior to onset of mortality, the differences cannot explain the increased mortality observed in the WD compared to low fat diets. Our data support a potential role for TcdA in the increased mortality with the WD, but via a mechanism independent of intestinal inflammation.

**Figure 2:**
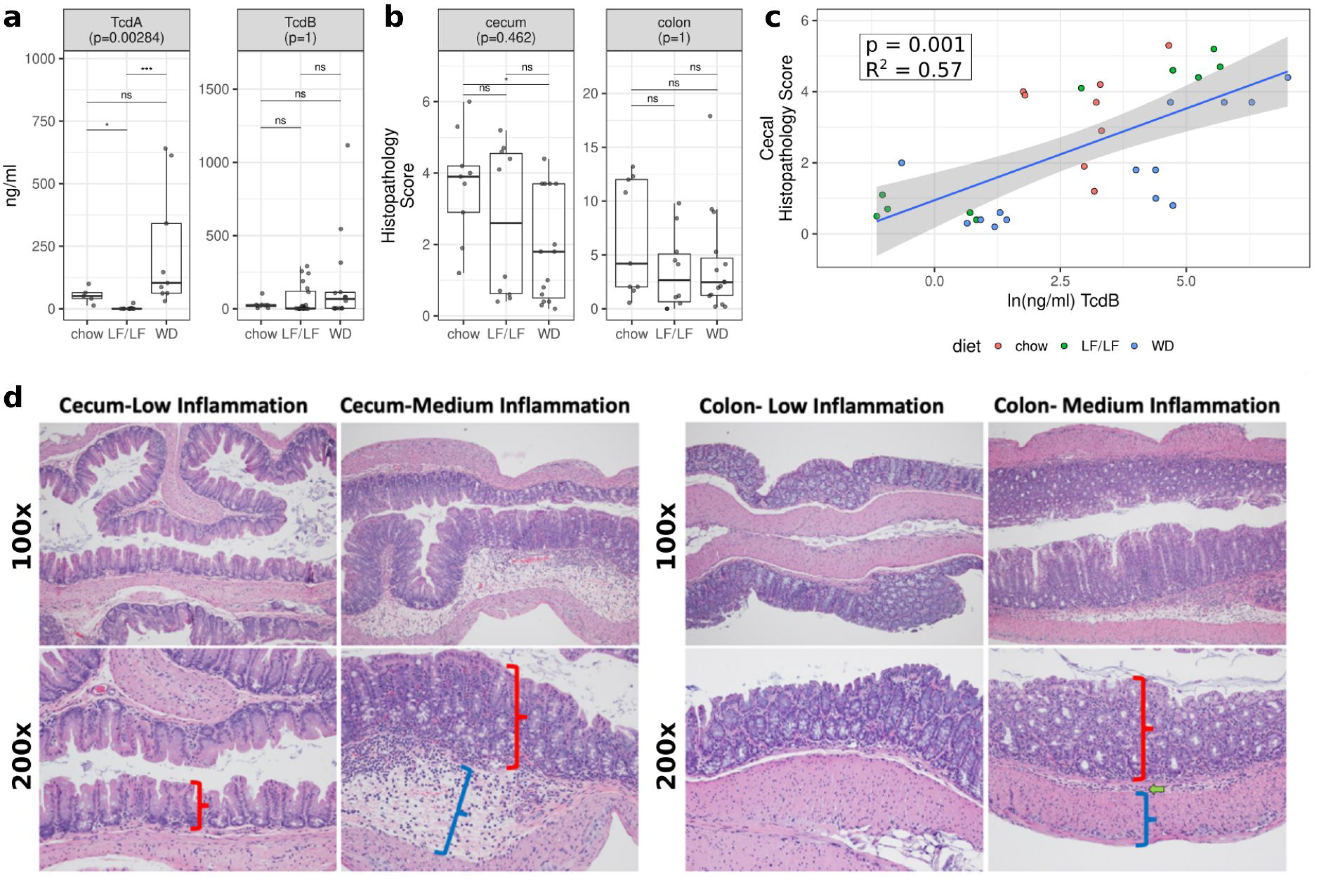
Toxin and histopathology scores by diet. **(A)** TcdA and TcdB production across diets as determined by ELISA. **(B)** Histologic inflammation scores across diets as determined by a blinded histologist. Significant differences were calculated with a Kruskal-Wallis and Dunn‟s post hoc test. Kruskal-Wallis p-values were corrected for multiple comparisons with the FDR algorithm of Benjamini and Hochberg ^25^. Also see Figure S2 for statistical analysis with linear modeling that controlled for batch. Median and Interquartile Range (IQR) indicated. (* : p < 0.05, ** : p < 0.01, *** : p < 0.001, **** : p < 0.0001) **(C)** Linear regression of cecal histopathology against *C. difficile* TcdB burden. (p = 0.001 with model cecum_infl ∼ TcdB and FDR correction). (D) Example sections of cecal (left) and colon (right) tissues with low or medium inflammation. No samples had levels of inflammation considered to be high. The cecum was scored for injury according to the system of Barthel et al, 2003 ^24^. Scoring of inflammation using the Barthel scoring system is restricted to neutrophils in the mucosa portion of the cecum. In low cecal inflammation, no neutrophils are observed in the mucosa (red bracket) while they are observed with medium inflammation. Medium cecal inflammation also displayed submucosal edema (blue bracket) that is thought to occur at least to some degree due to the neutrophils present in the submucosa. The colon (right panels) was scored for injury according to the system of Dieleman et al, 1998 ^23^. This system takes into account the relative quantity of inflammatory cells as well as whether they are found only in the mucosa layer (red bracket), are also in the submucosa (green arrow), or are found all the way through the muscularis (blue bracket) and into the peritoneal cavity.

### Cecal levels of bile acids and their relationship to diet, cecal levels of C. difficile toxins, and inflammation

To further explore mechanism, we used targeted LC/MS to measure the levels of a pool of 13 different bile acids in the aspirated cecal contents of a separate cohort of mice that were sacrificed at 3 days post infection (Table S2). Bile acids have a complex relationship with *C. difficile* germination and growth^13, 26–29^. The primary bile acids TCA and CA can promote the germination of *C. difficile* spores *in vitro*^13^ and primary bile acids including CA are elevated in individuals with first time or rCDI compared to controls ^30, 31^. The primary bile acid chenodeoxycholic acid (CDCA) can block TCA-induced spore germination^27, 28^ and another primary bile acid, Ursodeoxycholic acid (UDCA), can inhibit both *C. difficile* spore germination and *C. difficile* growth ^29^. Furthermore, the murine primary bile acids alpha muricholic acid (a_MCA) and beta muricholic acid (b_MCA) can inhibit *C. difficile* spore germination and growth ^32^. Of particular interest in this study are also the secondary bile acids DCA and lithocholate (LCA); these molecules are produced by the metabolic transformation of primary bile acids by intestinal microbes ^14^, can arrest the growth of vegetative *C. difficile* ^13^, and are lower in individuals with CDI ^30, 31^. We measured the levels of these bile acids with known effects on *C. difficile* as well as 5 other taurine-conjugated bile acids (Figure S3). To consider known effects of bile acids on *C. difficile* growth and germination in our analyses, we binned the bile acids that were inhibitors of *C. difficile* germination and/or growth (*CDCA, UDCA, a_MCA, b_MCA, LCA, DCA)*, and *C. difficile* germination promoters (*TCA* and *CA*). We also evaluated the ratio of *C. difficile* promoters to inhibitors, as has been done previously ^19^.

When comparing across diets, we were most interested in bile acid measures that showed differential levels between the WD and both the LF/LF and chow diets, since the WD had high CDI mortality compared to both the LF/LF and chow diets. *C. difficile* inhibitors were significantly lower in the WD compared to the chow diet, but there was not a difference between the WD and the LF/LF diet (Figure 3A). *C. difficile* promoters had significantly higher levels in the WD compared to chow but not compared to the LF/LF diet (Figure 3A). Interestingly, the ratio of promoters:inhibitors was significantly higher in the WD compared to both the LF/LF diet and chow, consistent with mortality differences. We also analyzed whether each of the 13 bile acids individually differed across diets, and diet significantly affected the levels of most (Figure S3). However, none were individually significantly different in the WD compared to both the LF/LF and chow diet. While regressing cecal and colon inflammation scores (Figure 3B) and toxins TcdA and TcdB (data not shown) against bile acid summary measures, the only significant relationship observed was a negative correlation between *C. difficile* inhibitors and colon inflammation. Evaluating differences across diets using linear regression models that included batch did not affect the interpretation of these results (Fig S2). Taken together, these data support that bile acid pools were strongly influenced by diet, with the WD having the most pro-*C. difficile* bile acid composition, but more work needs to be done to determine the degree to which these differences were driving the higher mortality observed with the WD.

**Figure 3:**
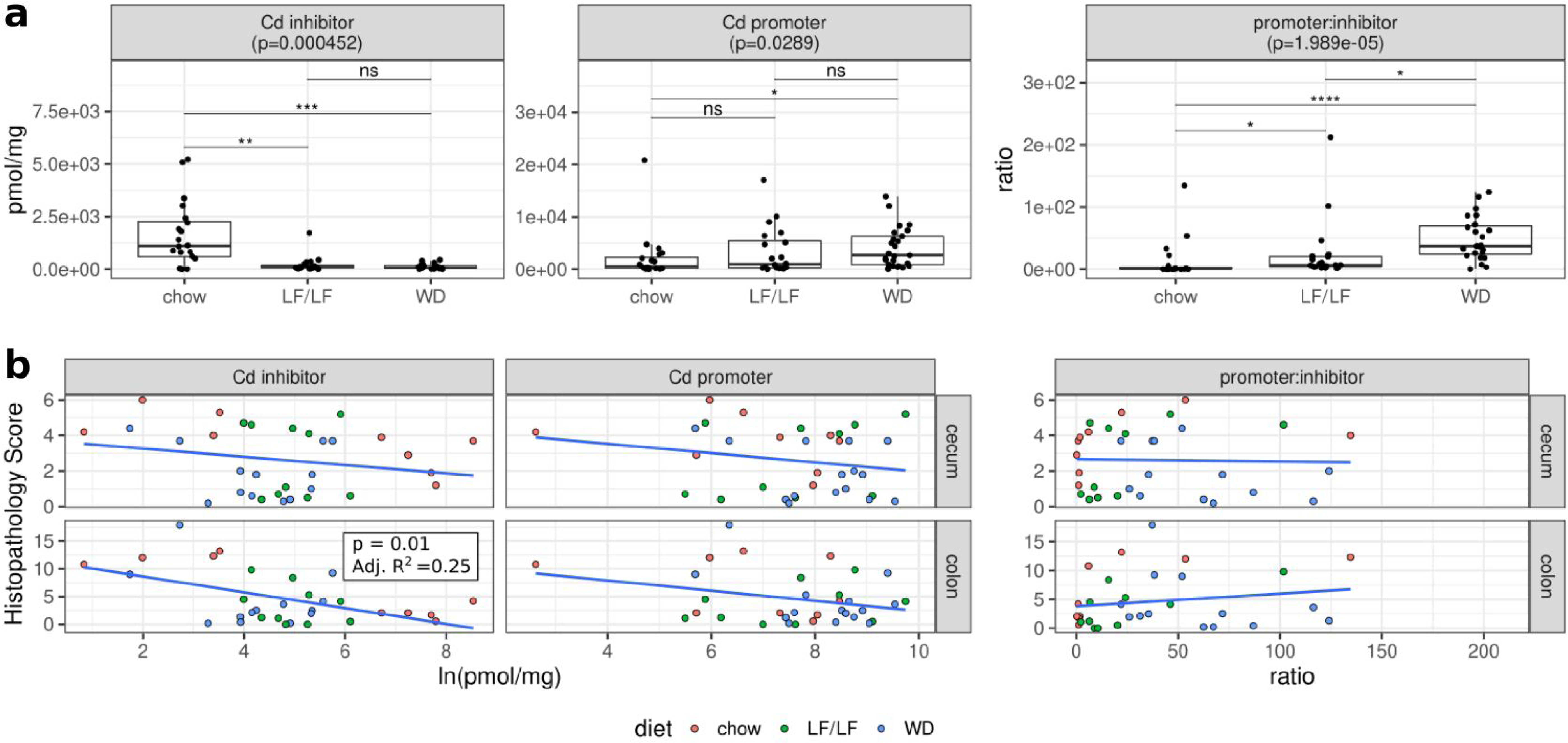
Bile acid pools in cecal contents of infected mice 3 days post C. difficile infection. **(A)** Cecal levels of *C. difficile* inhibitors (*CDCA, UDCA, a_MCA, b_MCA, LCA, DCA)*, *C. difficile* promoters (*TCA*, *CA*), DCA, and ratios of promoters:inhibitors across diets (chow n = 20, LF n = 20, WD n = 25; Table S2). Significant differences calculated with a Kruskal-Wallis and Dunn‟s post hoc test. Kruskal-Wallis p-values were corrected for multiple comparisons with the FDR algorithm of Benjamini and Hochberg. Median and IQR indicated. (* : p < 0.05, ** : p < 0.01, *** : p < 0.001, **** : p < 0.0001) **(B)** Linear regressions of cecal or colonic histology against CD inhibitors, CD promoters, and the promoter:inhibitor ratios. Only colon inflammation versus *C. difficile* inhibitors was significant.

### Cecal levels of SCFAs and their relationship to diet, secondary bile acids, cecal levels of C. difficile toxins, and inflammation

To further explore potential mechanisms of increased mortality in the WD-fed mice we also used targeted GC/MS to measure the levels of the SCFAs butyrate, propionate, and acetate in the aspirated cecal contents in mice that were sacrificed at 3 days post infection (Table S2). SCFAs are of interest because they are microbial products of fermentation of dietary fiber, have been previously implicated in the positive effects of a diet rich in MACs, ^15^ and are decreased in individuals with rCDI ^16^. Although butyrate can directly suppress *C. difficile* growth *in vitro* ^15^, butyrate also enhances *C. difficile* toxin production *in vitro* ^15^ ^33^. We also directly evaluated the secondary bile acid DCA, since it can arrest the growth of vegetative *C. difficile* ^13^ and prior studies have shown that secondary bile acid producers such as *Clostridium scindens* can protect against CDI in mice^14^.

Butyrate, acetate, and DCA were all significantly higher in the chow diet compared to both the LF/LF diet and WD (Figures 3A and 4A). There was also a significant correlation between levels of DCA and butyrate in a multivariate regression that accounted for differences across diets (Figure 4B). This is consistent with both DCA and butyrate having been linked with the presence of a healthy protective gut microbiome composition and low levels of both have been observed in individuals with rCDI^30, 34^. Surprisingly, butyrate positively correlated with TcdB (Figure 4C) and cecal and colonic inflammation (Figure 4D), but linear regression indicated that this relationship was dependent on diet, being driven by a positive association in the WD and LF/LF diet contexts only (Figure 4C, Table S3). DCA also correlated with TcdB levels and cecal and colonic inflammation in a diet dependent manner, with a positive relationship in LF/LF and WD and the expected negative (protective) relationship only in chow (Figure 4D, Table S3).

**Figure 4:**
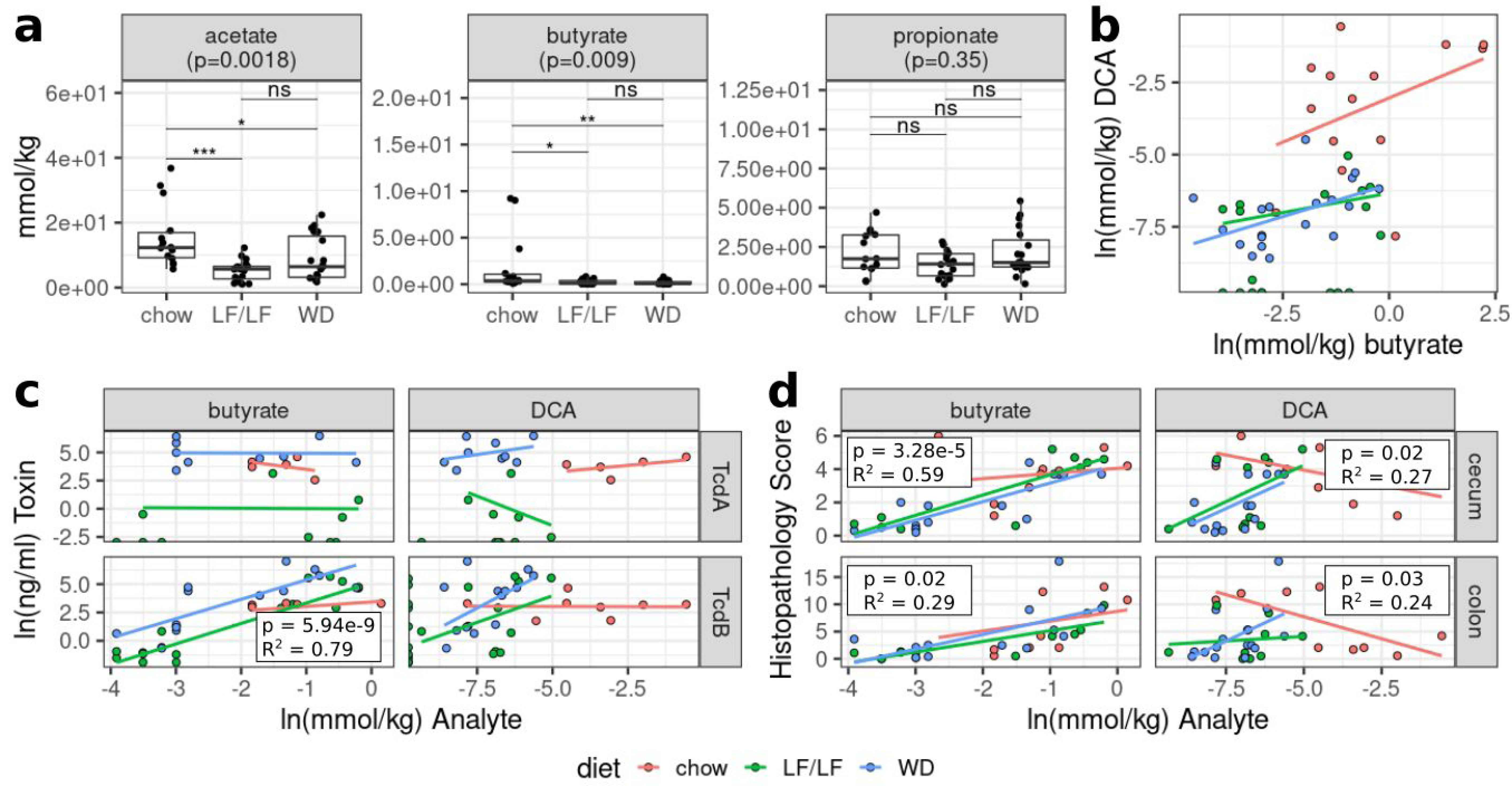
Relationships between microbial metabolites (SCFAs and the secondary bile acid DCA) and diet, toxin, and inflammation. **(A)** Cecal levels of the SCFAs acetate, butyrate, and propionate. p-values were determined using a Kruskal-Wallis with Dunn‟s post hoc test. Median and IQR indicated. (* : p < 0.05, ** : p < 0.01, *** : p < 0.001) **(B)** Multiple linear regression of DCA levels as a function of butyrate and diet. Model = ln(DCA) ∼ ln(butyrate) + diet + ln(butyrate)*diet. R-squared 0.855 p <0.0001. **(C)** Multiple linear regressions of *C. difficile* toxin TcdA and TcdB concentrations against butyrate and DCA while controlling for dietary interactions. **(D)** Multiple linear regressions of cecal or colonic histology against butyrate and DCA while controlling for dietary interactions.

### A conventional chow diet increases homogeneity of response, resilience and alpha-diversity of the gut microbiome after challenge with antibiotics and CDI compared to both purified diets

We next sought to understand how the composition of the fecal microbiome was affected by diet during the course of antibiotic treatment and infection with *C. difficile* (Fig. 1A). Fecal pellets were collected during experiment 1 upon arrival prior to diet change (Day -7), just prior to the start of oral antibiotic delivery (Day 0), after 5 days of oral antibiotics (Day 5), and daily through Day 10, which captured before and after the clindamycin injection given on day 7 and *C. difficile* gavage on Day 8 (Fig. 1A). Collected samples were subjected to 16S ribosomal RNA (rRNA) gene amplicon sequencing targeting the V4 region of the rRNA gene on the MiSeq platform.

Principle coordinate analysis (PCoA) plots of a weighted UniFrac ^35^ distance matrix suggested that mice fed either the WD or LF/LF diet had decreased resilience and a less homogeneous response to antibiotic challenge and CDI as compared to chow-fed mice (Figure 5). Mice fed either the WD or LF/LF diet showed greater divergence across PC1 upon antibiotic exposure than chow-fed mice, higher spread across mice in the same diet group, and less recovery towards their baseline after antibiotics (Fig. 5A). We quantified resilience by comparing the pairwise weighted UniFrac distances of mice across the experiment to baseline microbiota of their respective diet cohort at Day 0 (7 days post-diet change and pre-oral antibiotics; Fig. 5B). Chow-fed mice had significantly smaller weighted UniFrac distances from their baselines than the other groups at Day 5 (post 5 days antibiotic challenge) that persisted through Day 10 despite some convergence after clindamycin injection (Day 8) (Fig. 5B). By Day 9, chow-fed mice again displayed higher microbiome resilience than both the WD and LF/LF diet groups. We also assessed the homogeneity of response to a disturbance among mice in the same diet group. As an example, low homogeneity would occur if the mice within a diet group showed high variability in the degree to which their gut microbiome changed upon antibiotic exposure. We quantified this as the median pairwise weighted UniFrac distance for comparisons within samples collected at the same time point from mice fed the same diet (Fig. 5C). Both the WD and LF/LF diet showed much lower homogeneity of gut microbiome compositional response to antibiotic challenge, particularly to the 5-day treatment with oral antibiotics (Day 5), compared to chow-fed mice (Fig. 5C).

**Figure 5:**
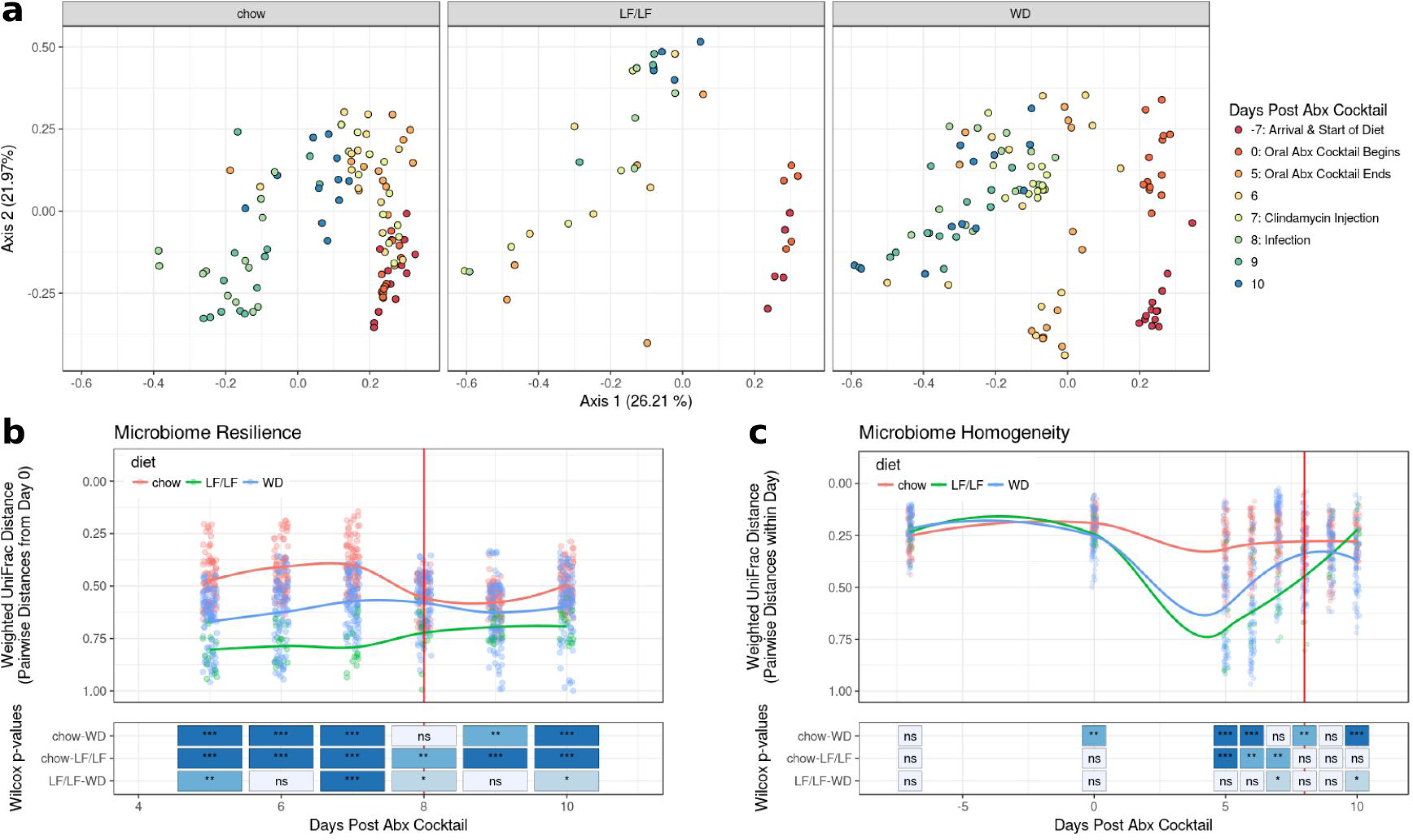
Beta diversity plots of fecal microbiome by diet during antibiotic treatment and infection with C. difficile. Vertical red lines in panels B and C designate the day of *C. difficile* infection (chow n = 13, LF n = 5, WD n = 13). **(A)** Weighted UniFrac PCoA plots of all samples with each diet highlighted in separate panels. **(B)** Resilience of microbiome composition assessed by within-mouse pairwise weighted UniFrac distances between Day 0 (7 days post diet switch and prior to oral antibiotics) and later time points and **(C)** Longitudinal plot of microbiome turnover homogeneity as plotted by intra-time point pairwise Weighted UniFrac distances within diet groups. Significant differences between diet groups were calculated by Kruskal Wallis followed by Dunn‟s post hoc test. Trend lines were fit using local polynomial regression. ***: p<0.001. **: p<0.01, *: p<0.05 ns= non-significant.

Similar patterns were seen when evaluating changes in alpha-diversity across the experiment between each diet cohort. Figure 6 shows changes in phylogenetic entropy, which is a measure of alpha diversity that considers species richness, evenness, and distinctness^36^. The phylogenetic entropy of the WD-fed mice was lower than chow-fed mice after diet change and this difference became more pronounced upon oral antibiotics and remained so through the rest of the experimental timeline (Fig. 6). Interestingly, the phylogenetic entropy of the LF/LF diet-fed mice remained equivalent to the chow-fed cohort with diet change but decreased to the same level as the WD with antibiotic treatment (Fig. 6).

**Figure 6:**
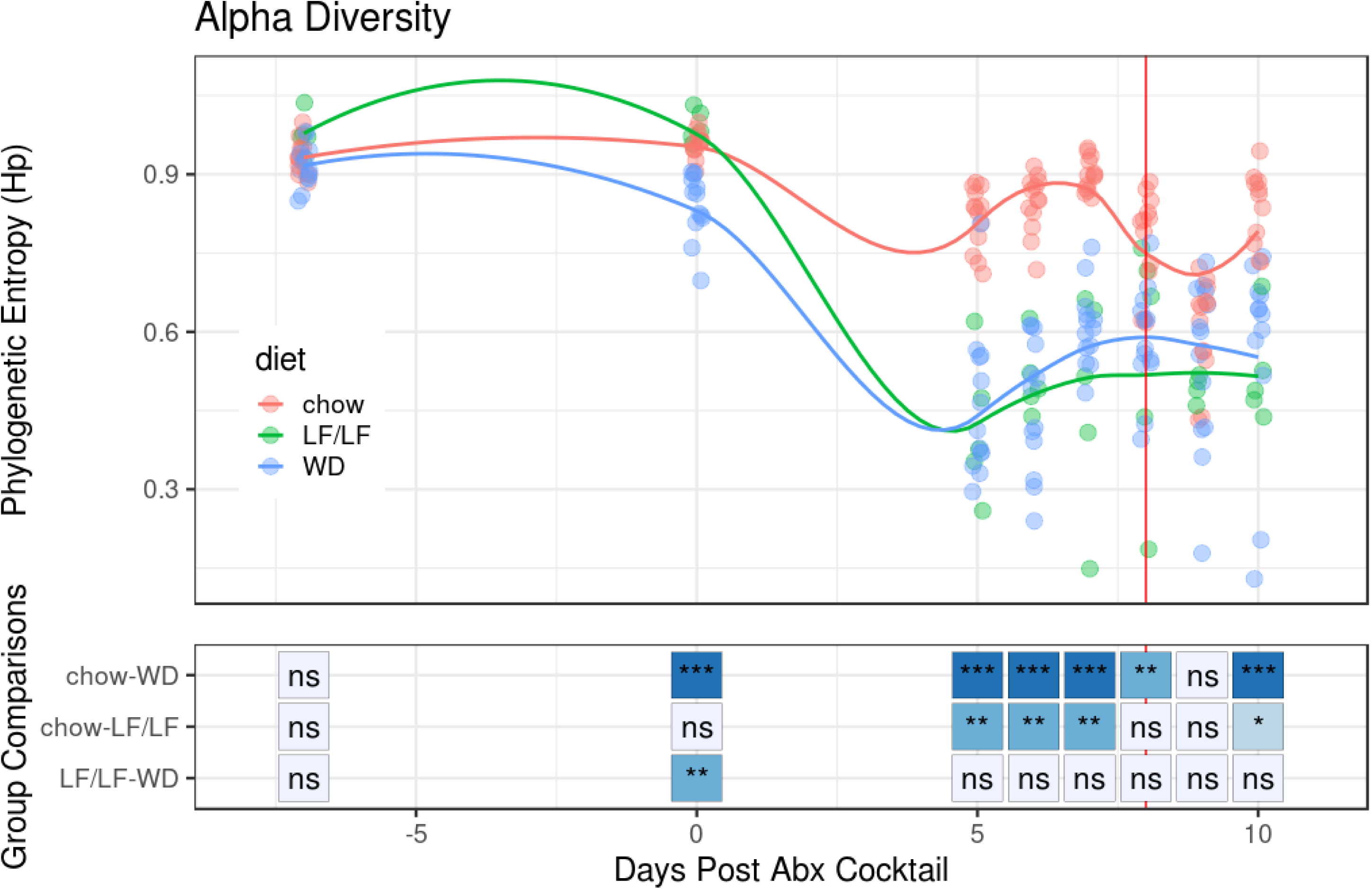
Alpha-diversity (phylogenetic entropy) of the fecal microbiome during murine CDI model. Data for each individual mouse is plotted as well as the fitted local polynomial regression for each diet group. Significant differences between diet groups were calculated by Kruskal Wallis followed by Dunn‟s post hoc test. * : p < 0.05, ** : p < 0.01, *** : p < 0.001.

### The WD and LF/LF diets had increased facultative anaerobe colonization and decreased secondary bile acid and SCFA-producing bacteria compared to conventional chow diet

Low-diversity dysbiosis is a state of disturbance that is often characterized not only by low alpha-diversity, but also by an increased ratio of facultative to strict anaerobes ^37^. Low-diversity dysbiosis is associated with a number of diseases including rCDI ^37^. We sought to investigate whether the different diets tested influenced if the microbiome developed a compositional state characterized by high levels of facultative anaerobe colonization and lower levels of strict anaerobes. Since Lactobacillales and Enterobacterales contain many important intestinal facultative anaerobes and most members of Clostridiales are strict anaerobes and include key butyrate and secondary bile acid producers, we plotted the relative abundances of these orders over the course of the experiment (Fig. 7A). All mice had decreases in the relative abundance of Clostridiales in their fecal microbiome with oral antibiotics; however, mice fed a chow diet were able to maintain a Clostridiales population while both the WD and LF/LF diets saw near-complete elimination of these taxa (chow-WD p<0.01 for days 0 through 9 and p<0.05 on day 10, Fig. S4). Conversely, mice fed either the WD or LF/LF diet had a large bloom of Lactobacillales after oral antibiotic treatment that was not observed in the chow-fed mice (chow- WD p <0.001 and chow-LF/LF p<0.05). Lastly, all 3 diet groups had a large increase in Enterobacterales in their fecal microbiome following antibiotics; however, the LF/LF and chow groups showed earlier decrease than WD mice (chow-WD p <0.01 and p<0.05 at days 9 and 10 respectively, Fig. S4). Comparisons of the LF/LF diet were limited due to smaller sample size (n = 5 vs. n = 13 for chow and WD).

**Figure 7:**
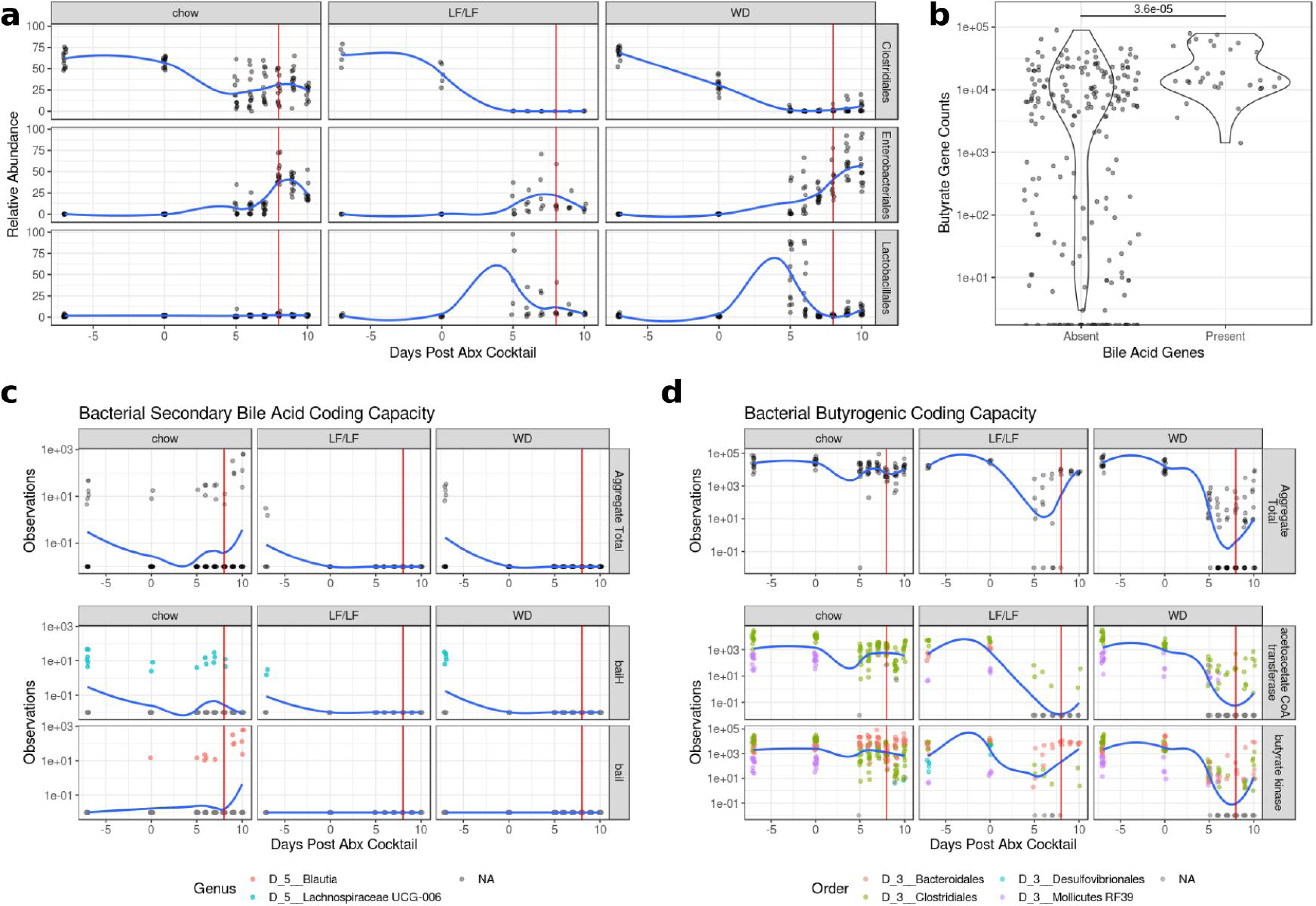
Changes in key taxa, and secondary bile acid and butyrate coding capacity during the CDI protocol. The vertical red line in (A), (C) and (D) indicates the day of *C. difficile* infection. All trend lines were fit using local polynomial regression. **(A)** Relative abundance of key bacterial orders during antibiotic treatment and infection. A summary of significant differences of these taxa across diets are in Fig. S4. **(B)** Violin plots of abundance of butyrate genes from PICRUSt2 analysis binned by presence of secondary bile acid producing genes (Wilcoxon p<0.001). **(C)** Time course of coding capacity of secondary bile acid genes. The top row shows the total capacity of each sample (*baiH* and *baiI*) while the bottom two rows show specific taxa contributions of key genes in the Bai operon. **(D)** Time course of coding capacity of butyrate producing genes by diet. The top row shows the total capacity as measured by *but* and *buk* genes while the bottom two rows show specific taxa contributions of *but* and *buk* specifically. Taxa with mean relative abundance < 0.01% were filtered from the analysis.

We also used PICRUSt^38^ to predict metagenomes using our 16S rRNA data to 1) investigate trends in the prevalence of key genes in secondary bile and butyrate production over the course of our experimental timeline and 2) predict which bacterial taxa were contributing these genes. Because *baiA*, *baiB*, and *baiCD* are not available in PICRUSt2‟s set of predicted genes, we only used the genes for *baiH* (KEGG ID: K15873) and *baiI* (KEGG ID; K15874), which are both genes in the *bai* operon^39^, to assess genomic potential for secondary bile acid metabolism. Acetoacetate co-A transferase (*but*; K01034) and Butyrate Kinase (*buk*; KEGG ID: K00929), which are the main pathways for fermentative production of butyrate in the gut microbiome ^40^, were used to assess butyrate production potential. Plotting these genes/pathways over time reveals a significant effect of diet on their abundance and response to antibiotics (Fig. 7C). Although all diet groups showed a marked decrease in bile acid genes with oral antibiotics, only the chow-fed mice displayed a recovery of secondary bile acid genes, though the source of these genes switched from Lachnospiraceae UCG-006 to Blautia. This result is consistent with our observation of higher cecal levels of secondary bile acids in chow-fed mice compared to mice fed either the WD or LF/LF diets at 3 days post *C. difficile* gavage (Fig. 4A).

Butyrate coding capacity also differed between diet groups. Chow-fed mice showed minimal change in the abundance of both the *but* and *buk* genes for fermentative butyrate production during the time course while the WD mice had a decrease of 5 orders of magnitude (Fig. 7D). The LF/LF diet-fed mice showed an intermediate phenotype with the resilience of the butyrate pathway being mostly attributed to a butyrate kinase dependent pathway. The results for *but* and not *buk* however are consistent with our measurements of cecal butyrate levels in these mice 3 days post *C. difficile* gavage (Fig. 4A). This is consistent with *but* being regarded to be a more important source of butyrate in the intestine ^41^.

Since we had observed a strong positive correlation between cecal levels of butyrate and the secondary bile acid DCA in our mass spectrometry data (Fig. 4B), we also determined whether there was a relationship between butyrate and secondary bile acid coding capacity. We found a highly significant association (p = 3.6x10^-5^), with secondary bile acid producing genes only predicted to be present in samples that also had high predicted levels of butyrate producing genes (Fig. 7B).

## Discussion

*C. difficile* infection is a grave and growing health threat. Current strategies to limit its spread have focused on sanitation and antibiotic stewardship, however incidence has continued to rise despite these efforts, highlighting the need for new treatment and prevention strategies ^2^. Because of the ubiquity of *C. difficile* spores in the environment and high levels of colonization, focusing on ways to increase the resilience of the host to *C. difficile* disease is one important prevention strategy ^42^.

Our results augment a growing body of evidence from studies conducted in mouse models of antibiotic induced *C. difficile* infection that points to dietary intervention as a promising approach to prevent antibiotic-induced CDI ^4, 15, 18, 19^. Prior studies have suggested the importance of a variety of macronutrients, including MACs, protein, and fat. Specifically, for MACs, one study that used an antibiotic-induced murine model demonstrated that mice fed a diet deficient in MACs (e.g. soluble fiber, resistant starches) had persistent *C. difficile* shedding and that there was a resolution of colonization with the reintroduction of inulin or other MACs ^15^. A protective effect of dietary MACS was also demonstrated in a human clinical trial in which a decrease in *C. difficile* recurrence from 34.3% to 8.3% was observed with prebiotic supplementation ^43^. Our results are consistent with these studies in that both the WD and LF/LF diets were low in MACs compared to the chow diet and had a greater antibiotic-induced disturbance to the gut microbiome and loss of CDI protective microbial metabolites such as DCA. However, it is important to note that the differences between conventional chow, WD and LF/LF diets extend well beyond fiber, and these other dietary components could also have influenced our observation. Also, our LF/LF diet had greatly reduced mortality compared to the WD, even though the LF/LF diet-fed mice was low in MACs and had a comparable level of microbiome disturbance and loss of protective metabolites with antibiotic treatment.

Influence of dietary protein has also been noted in a few studies. Specifically, one study found a low-protein diet to be protective in an antibiotic-induced CDI murine model, with mice fed a 2% protein diet having increased survival, decreased weight loss, and decreased overall disease severity compared to mice fed a 20% protein defined diet ^20^. Another study showed that a diet poor in proline (an essential amino acid for *C. difficile* growth) prevented *C. difficile* carriage ^4^. Furthermore, in a recent study that evaluated both a high-fat/high-protein Atkins-type diet and a high-fat/low-protein diet in a mouse model of antibiotic-induced CDI, the high-fat/high-protein diet promoted severe CDI and 100% mortality, while the high-fat/low-protein diet had variable disease severity and survival, showing a strong effect of dietary protein but indicating that the effects of fats were uncertain ^18^.

Another had found that a diet that was high in refined carbohydrates and low in fiber had improved CDI severity compared to mice fed a standard chow diet ^18^. New data has suggested that novel speciation of *C. difficile* may be selecting for strains that show increased sporulation and host colonization capacity with sugar availability (glucose or fructose) ^47^. This work, conducted with *C. difficile* strain (VPI 10463), did not show differences in mortality from CDI in low-fat/low-fiber diets with different amounts of sucrose ^18, 19^.

Our results show that high dietary fat in the context of low dietary fiber had a strong effect on CDI-induced mortality, with mechanisms distinct from a loss of beneficial microbial metabolites. Evidence to suggest that a high-fat/low-fiber western-type diet could have a profound effect on CDI was first presented over 20 years ago in experiments designed to study the atherogenic properties of a Western diet in Syrian hamsters ^44, 45^. Significant mortality from CDI was observed in hamsters fed a high-fat/low-fiber pro-atherogenic diet and not a typical high-fiber/low-fat hamster diet, even in the absence of an antibiotic disturbance ^44, 45^. Another recent study that conducted a study of antibiotic-induced CDI in a high-fat-diet (HFD) induced obesity model found protracted disease in the HFD compared to a chow diet ^19^, but not the severe mortality that we observed with a high-fat/low-fiber diet. We posit that high dietary fat may have a more profound influence on CDI than low dietary fiber since a prior study of MAC deficient diets found that low fiber was associated with higher *C. difficile* carriage but did not describe the severe disease/mortality that was observed here while using a similar mouse model ^15^. However, since we did not test a high-fat/high-fiber diet, it is unclear whether the high mortality that we observed was due to a combination of high-fat and low-fiber in the diet, or just dietary fat.

Although these studies taken together support a potential synergy of high-fat and low-fiber leading to severe disease, it is important to note that these papers differ in many experimental parameters including the source of the mice (which has been shown to influence response to antibiotic perturbation and *C. difficile* clearance in mice ^46^), types of antibiotics used, strain of *C. difficile*, and whether *C. difficile* was used as active growing bacteria (as done in our study) or as spores.

### The Role of Toxin Production and Inflammation

In order to further explore potential causes of death, we looked at both inflammation by histology and levels of the toxins TcdA and TcdB by ELISA in cecal contents collected 3 days post *C. difficile* infection, which was just prior to the onset of mortality in our longitudinal cohort. Both TcdA and TcdB can disrupt cytoskeletal structure and tight junctions of target cells ^48^ and induce inflammation ^3, 49^. We did not observe any differences in TcdB or cecal or colon inflammation scores across diets. However, cecal levels of TcdB did correlate with cecal inflammation, consistent with known effects of TcdB ^48^ ^50^. This supports that levels of TcdB produced by *C. difficile* may indeed be causing pathology in these mice, but higher levels of TcdB at Day 3 post CDI cannot alone explain the higher mortality that we began to observe at Day 4 post CDI infection in the WD-fed mice. Interestingly, TcdA and TcdB levels did not correlate with each other, and TcdA levels did show a pattern at Day 3 post-CDI consistent with mortality, being significantly higher in the WD mice compared to the LF/LF-fed mice. Consistent with the LF/LF diet mice having delayed mortality compared to chow-fed mice, there were also lower levels of TcdA in the LF/LF fed mice compared to those fed a conventional chow diet at Day 3 post-CDI. However, unlike TcdB, TcdA levels did not correlate with inflammation. The lack of correlation of TcdA and TcdB with each other is surprising since they are often co-expressed, although their transcription is regulated by unique promoter regions^51^, and other post-transcriptional factors at the RNA or protein degradation level may also be at play. Studies of the activity of TcdA versus TcdB in various animal models have more strongly supported the importance of TcdB in CDI pathogenesis, and studies investigating TcdA have had mixed results^48^, although none of these studies were conducted in the context of a high-fat/low-fiber diet. Our results support a potential importance of TcdA and not TcdB in diet-associated differences in CDI pathogenesis, but further studies that sample the toxin levels at more time points over disease progression might prove illuminating. Indeed, another prior study that showed higher CDI pathology in HFD-induced obesity model versus a regular chow diet did not observe higher toxin levels (while binning TcdA and TcdB ELISA data) at day 3 post infection (acute phase), but did find higher toxin levels and intestinal inflammation between diets at day 10 post infection, due to recovery occurring in the chow fed but not HFD-obese mice ^19^.

Although it is possible that differences in TcdB and inflammation across diets in our study may have emerged over time, it was not possible to evaluate this since we had much higher mortality in our model, and most of our WD fed mice would have died by day 10. Further studies that use complementary methods to measure toxin besides just ELISA, which can lack specificity for TcdB in particular^52, 53^, or with strains of *C. difficile* that produce TcdA or TcdB only would be required for further validation ^52^. Also, in these studies we measured toxin levels but were unable to produce quality data regarding levels of *C. difficile* bacteria in the cecal materials. We thus cannot evaluate whether these differences in toxin levels are driven by more bacteria or increased toxin production by similar loads of bacteria.

### Effects of bile acids

Differences in host bile acid production and microbial bile acid metabolism is one potential mechanism of high-fat diet induced modulation of CDI severity. In the clinical setting, studies have shown that patients with CDI have increased TCA and decreased concentrations of the secondary bile acids DCA and LCA in their feces ^30, 34^ as well as other complex alterations to bile acid pools ^31^. These derangements are corrected with fecal microbiota transplant for treatment of *C. difficile* (FMT) ^34^. *In vitro* experiments have shown that the primary bile acids TCA and CA are potent *C. difficile* germination factors ^13^ while UDCA and CDCA have been shown to inhibit germination and growth of *C. difficile in vitro* ^27–29^. The microbially-produced secondary bile acids DCA and LCA have also been shown to affect *C. difficile in vitro*: DCA promotes germination of *C. difficile* spores ^13^ while LCA inhibits germination ^29^ and both inhibit growth of vegetative *C. difficile* ^13, 29^. In line with these effects, reduced prevalence of the secondary bile acid producer, *Clostridium scindens* in the fecal microbiome has been associated with high incidence of CDI in both humans and in experimental mouse models, and gavaging mice with *C. scindens* protected against CDI and restored intestinal secondary bile acid levels ^14^. Despite this strong evidence of a role of a protective effect of microbially produced secondary bile metabolites in protection from CDI, this mechanism did not appear to be a sole driving factor of the mortality that we observed in mice fed a WD, since the levels of these metabolites were lowest in the mice fed the LF/LF diet even though the LF/LF mice did not experience increased mortality. Levels of *C. difficile* inhibitors, which included the secondary bile acids DCA and LCA, did negatively correlate with colonic inflammation, suggesting some degree of protection in these mice. Functional interrogation of the microbiome using PICRUSt suggests that the lack of secondary bile acids in the WD and LF/LF diet fed mice might be due to a lack of recovery of secondary bile acid producing bacteria following antibiotic disturbance in both the WD and LF/LF diet contexts.

We did find that the ratio of *C. difficile* promoters:inhibitors was significantly higher in the WD compared to both the LF/LF and chow diets, consistent with mortality differences. Our results support that a high-fat diet coupled with low-fiber and antibiotic treatment may provide a “double hit” for shifting towards a pro-*C. difficile* bile acid pool – with dietary fat increasing excretion of pro-*C. difficile* primary bile acids into the gut and antibiotic-induced gut microbiome disturbance decreasing their conversion into protective secondary bile acids. *In vitro* assays have demonstrated that variable mixtures of primary and secondary bile assays have different impacts on *C. difficile* germination and growth ^54^. However, more work needs to be done to determine the degree to which these differences were driving the higher mortality observed with the WD. A more convincing result would be if the *C. difficile* promoter:inhibitor ratio also predicted *C. difficile* toxin production while controlling for diet, but this was not the case (Fig. 3B).

We also found that diet had a significant effect on 4 of the 5 taurine conjugated bile acids that we assayed, with TCA, T_b_MCA, TDCA, and TCDCA all showing a pattern of increased levels in the WD compared to both the chow and LF/LF diets, but only comparisons of chow versus WD reaching statistical significance (Fig. S3). It is probable that the further decrease in levels of these taurine conjugated bile acids in the chow compared to the LF/LF diet is because the primary bile acids that are produced by the host are converted by microbes to secondary bile acids in only the chow diet. Our finding of increased TCA in the WD compared to chow is consistent with a prior study that found that IL10-deficient mice fed a diet high in saturated fat, had an increased proportion of taurine-conjugated bile acids compared to standard chow, and a diet high in poly-unsaturated fats ^21^. One prior study demonstrated that both TDCA and TCDCA have pro-germinative effects on *C. difficile*, though in our study, their cecal concentrations were orders of magnitude lower than TCA which is also a much stronger germinant ^55^.

One weakness of our study is that we cannot differentiate between the complex changes of the bile acid pools and the effects of the dietary components themselves – such as known effects of high-fat diet on inflammation ^56, 57^. Controlled studies that directly alter bile acid pools without also altering diet are valuable. In one study of CDI in HFD-induced obesity, inhibiting primary bile acid synthesis with the FXR antagonist obeticholic acid ameliorated CDI disease during later phases of infection but not in acute CDI ^19^. Another factor that may have influenced our result is that like other related studies ^15^, our infection procedure used a sample cultured for ∼24 hours without enumerating or enriching the sporulated fraction of the inoculum. As spores are the likely infective form of *C. difficile* in clinical settings, and many bile acids influence *C. difficile* pathogenesis by promoting or inhibiting germination, it is of interest to determine how the variability in vegetative composition influences the relationship between bile acid pools and CDI pathogenesis.

### Effects of the microbiome and their metabolites

Our data suggests that a complex diet is critical for the resilience and homogeneity of response of the gut microbiome after perturbation. In both cohorts of mice fed a purified diet that was deficient in fiber, the gut microbiome was significantly more variable and slower to recover to baseline after perturbation. We hypothesize that by supplying the gut with a preferred fuel (fiber) for species associated with health (e.g. strict anaerobes), the community is able to resist antibiotic induced changes and reconstitute more quickly once the pressure of antibiotic treatment has been removed. Since the chow diet differed from the purified diets in many components besides the levels of fiber, we cannot conclude from our study alone that increased resilience to microbiome disturbance with antibiotics in chow is driven by differences in fiber. However, our results are consistent with previous murine studies that have shown that low fiber diets can increase antibiotic-induced microbiome disturbance and delay recovery from treatment with ciprofloxacin ^58^ and that fiber supplementation can lead to a reduced disruption of the gut microbiome to disturbance from amoxicillin ^59^.

The increased resilience of gut microbiome composition to antibiotic disturbance was also reflected through levels of the bacterially produced metabolites that we measured. Neither the WD or LF/LF diets were able to maintain butyrate or secondary bile acid production following antibiotic perturbation. Based on the correlation between butyrate and DCA concentrations, we speculate that the lack of butyrate leads to increased luminal oxygen concentrations that are unsuitable for *Clostridium scindens* and other secondary bile acid producers. Prior work has shown that aerobic metabolism of butyrate by intestinal epithelial cells is a key driver of intestinal hypoxia ^60^. That there may be increased luminal oxygen concentrations in the LF/LF and WD is consistent with our observation of a bloom in Lactobacillales order, which is entirely composed of facultative anaerobes, after oral antibiotic challenge in the WD and LF/LF diets but not chow.

While our data do not suggest a role for fiber in protection against mortality from CDI in this mouse model since the LF/LF diet fed mice were protected without fiber in the diet, it would be short-sighted to dismiss the beneficial role of fiber in maintaining a healthy gut microbiome and resistance to CDI. Our model utilized a rather short-term diet change and an intense antibiotic regimen. We also did not explore diets high in fat and high in fiber, where it is possible that increased microbiome resilience to antibiotics due to fiber may protect from the detrimental effects of fat. As discussed above, a fiber-deficient diet has been shown to hinder clearance of *C. difficile* after challenge ^15^.

One surprising finding of our work, however, given these protective effects of dietary Microbiota Accessible Carbohydrates (MACs) from other studies, was that butyrate, a major fermentation product of MACs, positively correlated with TcdB and cecal and colonic inflammation, driven by an association in the WD and LF/LF diet contexts and not chow. DCA also correlated with butyrate and with TcdB levels and cecal and colonic inflammation in a diet dependent manner, with a positive relationship in LF/LF and WD and the expected negative (protective) relationship only in chow. Butyrate is typically associated with beneficial effects on gut health, including supporting intestinal barrier function ^61–63^, suppressing inflammation through induction of T regulatory cells ^64^, and directly suppressing *C. difficile* growth *in vitro* in a dose dependent manner ^15^. Also, lower butyrate and DCA has been observed clinically in individuals with CDI ^30, 34^. However, a positive correlation between butyrate and TcdB is consistent with prior studies showing that butyrate enhances *C. difficile* toxin production *in vitro* ^15, 33^, leading some to suggest that butyrate may signal to *C. difficile* a competitive gut environment ^15^. A similar diet-dependent detriment of SCFAs was observed in a study that showed that soluble fiber-supplementation drove hepatocellular carcinoma in mice in a manner dependent on microbial fermentation to SCFAs, but this effect occurred when soluble fiber was added to a compositionally defined diet and not to a conventional chow diet ^65^. Our results suggest that supplementation with soluble fibers such as inulin to prevent *C. difficile* may not produce the desired result in individuals who are otherwise consuming highly refined diets.

### Limitations of our study

We have demonstrated a striking difference in diet-mediated mortality in an antibiotic-induced murine CDI model, but our study does have limitations. We did not explore how the composition of fat influences these factors. Our WD composition represents a typical diet in the United States based on population survey data. Further studies to determine if total fat intake or specific types of fat drive our observed phenotype are needed. Furthermore, we only evaluated the effects of fat in a low-fiber context. Evaluating a high-fat/high-fiber diet would elucidate whether the expected beneficial effects of fiber on the microbiome would temper the negative effects of high-fat. Comparisons between chow-fed mice and those receiving a purified diet are limited due to the marked differences in the composition of macronutrients ^66^. Since this is an antibiotic-induced CDI model, our results only reflect effects of diet in the context of antibiotic disturbance. Finally, we note that we induced CDI infection using a standardized amount of live *C. difficile*, which is commonly used in murine studies of *C. difficile* ^22^. However, we note that since different bile acids influence spore germination as well as growth, results may vary in challenge models that instead use spores. For instance, we might expect the promotors TCA and CA to have a stronger effect in spore-infection models compared to our model that used gavage with vegetative forms of *C. difficile*, since they are potent germination factors of *C. difficile* spores. Future experiments to compare these results to a spore-infection model could elucidate the degree to which diet may affect CDI pathogenicity by influencing germination versus growth of *C. difficile* through modification of bile acid pools.

### Conclusions

This study along with recently published findings investigating dietary fiber ^15^, dietary proline ^4^, protein ^18^, and fat ^18, 19^ intake provides a compelling case that diet should be increasingly targeted as a prevention and treatment modality for CDI. High-risk populations such as elderly hospitalized individuals subjected to antibiotics and adult and pediatric oncology patients may benefit from decreased *C. difficile* colonization through diets with decreased fat and increased fiber. For patients with active infection, limiting fat intake could decrease disease severity while maintaining enteric nutrition.

## Methods

### Mouse diets

Diets were all obtained from Envigo (Indiana): Standard chow - Teklad global soy protein-free extruded (item 2920X - https://www.envigo.com/resources/data-sheets/2020x-datasheet-0915.pdf), Western Diet – New Total Western Diet (item TD.110919), Low-fat/low-fiber – variant of AIN93G (item TD.180811). See Table S4 for detailed composition of purified diets.

### Murine model of CDI

Mice were infected using a widely used murine CDI model ^22^ with minor modifications. Briefly, 6-week-old female C57BL/6 mice from Taconic Bioscience (Rensselaer, NY) arrived at University of Colorado on Day -7 of the experiment. During experiments, mice were cohoused in groups of 4-5 mice per cage. Survival experiments were conducted in four independent experiments at four separate starting dates. For cecal metabolite and toxin analysis, 2-7 independent experiments were conducted with separate starting dates (Table S2). Within 24 hours, mouse feed was changed to one of three diets: standard chow, high-fat/low-fiber (WD), or LF/LF diet (all groups n=20 over 4 batches; Table S2). After seven days of the new diet, we placed mice on a five-antibiotic cocktail (kanamycin (0.4 mg/ml), gentamicin (0.035 mg/ml), colistin (850 U/ml), metronidazole (0.215 mg/ml), and vancomycin (0.045 mg/ml)) in their drinking water. Antibiotics were removed for 48 hours, after which we administered an intraperitoneal injection of clindamycin in normal saline (10 mg/kg body weight). Twenty-four hours after injection, we gavaged mice with 1.75x10^5^ cfu of *C. difficile* VPI 10463 in the vegetative stage. We weighed mice daily after removal of oral antibiotics and they were euthanized if they lost >15% of body weight or were moribund. Fecal pellets were collected at arrival (Day -7), after diet change and prior to oral antibiotics (Day 0) and then daily after removal of oral antibiotics (Day 5-10). In a separate set of experiments, we performed the same experimental protocol on 66 mice (chow = 20, low-fat/low-fiber = 20, WD = 26) over 7 different batches (see Table S2), but we sacrificed the mice 72 hours after infection and collected cecal contents for SCFA, bile acid and toxin quantification and cecum and intestines for histopathology. Mice for the second experiments were also obtained from Taconic Bioscience. All mouse experiments were approved by the Institutional Animal Care and Use Committee and complied with their guidelines and NIH Guide for the Care and Use of Laboratory Animals (IACUC protocol #00249).

### C. difficile growth

*C. difficile* strain VPI 10463 (ATCC, Manassas Virginia) was used for all experiments. Frozen stocks were plated on to TCCFA agar plates (TekNova) and incubated overnight in an anaerobic chamber (Coy, Grass Lake, Michigan). Single colonies were picked and inoculated into BHI Media (Difco) and grown over night in anaerobic conditions. Cell quantities were quantified with flow cytometry using the BD Cell Viability Kit with BD Liquid Counting Beads (BD Biosciences). Cultures were then centrifuged at 3,000 g for 15 minutes and washed with sterile PBS three times before dilution into sterile water to a final concentration of 9x10^5^ cfu/mL.

### DNA Extraction and Sequencing

Total genomic DNA was extracted from fecal pellets from a subset of the mice in cohort 1 (chow = 13 mice from four separate cages over two experiments, WD = 13 mice from four separate cages over two experiments, LF/LF = 5 mice from two cages over two experiments) using the DNeasy PowerSoil Kit (Qiagen, Germantown, MD). Modifications to the standard protocol included a 10-minute incubation at 65°C immediately following the addition of the lysis buffer and the use of a bead mill homogenizer at 4.5 m/s for 1 min. The V4 variable region of the 16S rDNA gene was targeted for sequencing (515F: GTGCCAGCMGCCGCGGTAA, 806R: GGACTACHVGGGTWTCTAAT). The target DNA was amplified using 5Prime HotMaster Mix (Quantabio, Beverly, MA). Construction of primers and amplification procedures follow the Earth Microbiome Project guidelines (www.earthmicrobiome.org) ^67^. Amplified DNA was quantified in a PicoGreen (ThermoFisher Scientific) assay and equal quantities of DNA from each sample was pooled. The pooled DNA was sequenced using a V2 2x250 kit on the Illumina MiSeq platform (San Diego, CA) at the University of Colorado Anschutz Medical Campus Genomics and Microarray Core facility.

### Sequence Data Analysis

Raw paired-end FASTQ files were processed with QIIME 2 version 2018.8 ^68^. Denoising was performed with DADA2 ^69^, a phylogenetic tree was built using sepp ^70^ and taxonomy was assigned to amplicon sequence variants (ASVs) using the RDP Classifier ^71^ trained on the Silva version 132 taxonomic database ^72^ ^73^ using QIIME 2 ^68^. The data was rarefied at 5,746 sequences per sample. Alpha-diversity was measured by phylogenetic entropy ^36^ and beta-diversity was determined by weighted UniFrac distances ^35^. PCoA of weighted UniFrac plots were constructed using QIIME 2. Metagenomes were imputed from 16S ASVs using PICRUSt2‟s default pipeline for stratified genome contributions ^38^. Low abundance taxa (<0.01% mean relative abundance) were filtered for analysis of the butyrogenic coding capacity. Software was installed using Anaconda ^74^ and analysis was performed on the Fiji compute cluster at the University of Colorado Boulder BioFrontiers Institute.

### SCFA quantification

The SCFAs butyrate, propionate, and acetate were analyzed by stable isotope GC/MS as previously described ^75^. Briefly, cecal samples were collected directly into pre-weighed, sterile cryo vials and flash frozen at −80°C until processing. Samples were then subject to an alkylation procedure in which sample and alkylating reagent were added, vortexed for 1 min, and incubated at 60°C for 25 min. Following cooling and addition of n-hexane to allow for separation, 170 μL of the organic phase was transferred to an auto sampler vial and analyzed by GC/MS. Results were quantified in reference to the stable isotope standard and normalized to sample weight.

### Bile acids quantification

#### Reagents

LC/MS grade methanol, acetonitrile, and isopropanol were obtained from Fisher Scientific (Fairlawn, New Jersey). HPLC grade water was obtained from Burdick and Jackson (Morristown, New Jersey). Acetic acid, cholic acid (CA), chenodeoxycholic acid (CDCA), lithocholic acid (LCA), taurocholic acid (TCA) and deoxycholic acid (DCA) were obtained from Sigma Aldrich (St. Louis, Missouri). Taurodeoxycholic acid (TCDCA), taurochenodeoxycholic acid (TCDCA), taurolithocholic acid (TLCA), alpha-muricholic acid (a_MCA) and beta-muricholic acid (b_MCA) were obtained from Cayman Chemical (Ann Arbor, Michigan). Chenodeoxycholic acid-d4 (CDCA) and glycochenodeoxycholic acid-d4 were obtained from Cambridge Isotope labs (Tewksberry, Massachusetts).

#### Standards preparation

An internal standard containing 21 µM of chenodeoxycholic acid-d4 and 21 µM of glycochenodeoxycholic acid–d4 was prepared in 100% methanol. A combined stock of all bile acid standards was prepared at 0.5mM in 100% methanol. Calibration working standards were then prepared by diluting the combined stock over a range of 0.05 µM-50 µM in methanol. A 20 µL aliquot of each calibration working standard was added to 120 µL of methanol, 50 µL of water and 10 µL of internal standard (200 µL total) to create 10 calibration standards across a calibration range of 0.005 µM-5 µM.

#### Sample preparation

Fecal samples were prepared using the method described by Sarafian et al ^76^ with modifications. Briefly, 15-30mg of fecal sample were weighed in a tared microcentrifuge tube and the weight was recorded. 140 µL of methanol, 15-30 µL of water and 10 µL of internal standard were added. The sample was vortexed for 5 seconds, and then incubated in a -20°C freezer for 20 minutes. The sample was then centrifuged at 6000RPM for 15 minutes at 4°C. 185-200 µL of the supernatant was transferred to an RSA autosampler vial (Microsolv Technology Corporation, Leland, NC) for immediate analysis or frozen at -70°C until analysis.

#### High performance liquid chromatography/quadrupole time-of-flight mass spectrometry (HPLC/QTOF)

HPLC/QTOF mass spectrometry was performed using the method described by Sarafian et al ^76^ with modifications. Separation of bile acids was performed on a 1290 series HPLC from Agilent (Santa Clara, CA) using an Agilent SB-C18 2.1X100mm 1.8 µm column with a 2.1X5mm 1.8um guard column. Buffer A consisted of 90:10 water:acetonitrile with 1mM ammonium acetate adjusted to pH=4 with acetic acid, and buffer B consisted of 50:50 acetonitrile:isopropanol. 10 µL of the extracted sample was analyzed using the following gradient at a flow rate of 0.6mls/min: Starting composition=10% B, linear gradient from 10-35% B from 0.1-9.25 minutes, 35-85% B from 9.25-11.5 minutes at 0.65mls/min, 85-100% B from 11.5-11.8 minutes at 0.8mls/min, hold at 100% B from 11.8-12.4 minutes at 1.0ml/min, 100-55% B from 12.4-12.5 minutes 0.85mls/min, followed by re-equilibration at 10% B from 12.5-15 minutes. The column temperature was held at 60°C for the entire gradient.

Mass spectrometric analysis was performed on an Agilent 6520 quadrupole time of flight mass spectrometer in negative ionization mode. The drying gas was 300°C at a flow rate of 12mls/min. The nebulizer pressure was 30psi. The capillary voltage was 4000V. Fragmentor voltage was 200V. Spectra were acquired in the mass range of 50-1700m/z with a scan rate of 2 spectra/sec.

Retention time and m/z for each bile acid was determined by injecting authentic standards individually. All of the bile acids produced a prominent [M-H]^-^ ion with negative ionization. The observed retention time and m/z was then used to create a quantitation method. Calibration curves for each calibrated bile acid were constructed using Masshunter Quantitative Analysis software (Aligent Technologies). Bile acid results for feces in pmol/mg were then quantitated using the following calculation:

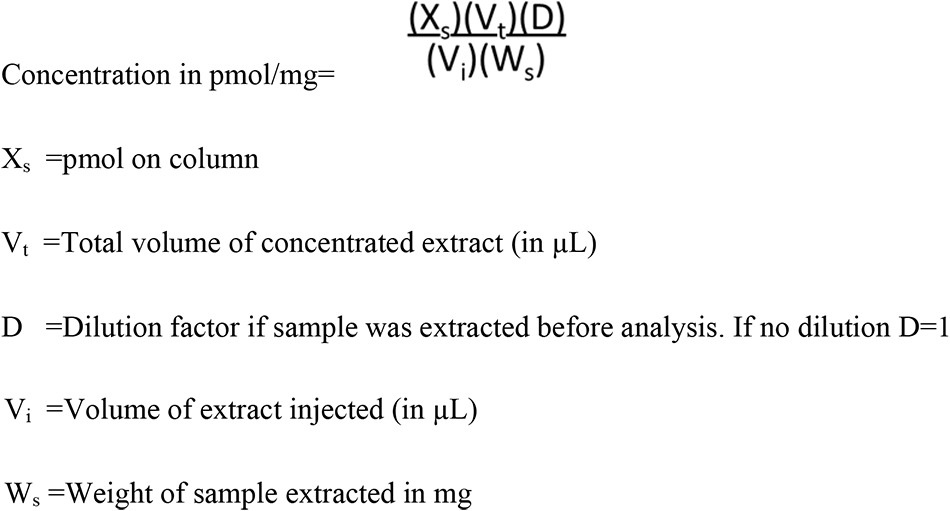

### *C. difficile* toxin TcdA and TcdB quantification

TcdA and TcdB concentrations were determined in cecal samples from day 3 of infection by comparison to a standard curve using ELISA (tgcBiomics, Germany). For samples that were too small to weigh accurately, a mass of 5 mg was assigned for concentration calculation. This mass was selected as it was the lowest weight that could be accurately determined.

### Histologic evaluation of large intestinal tissue

Cecum and transverse colon were harvested from mice three days after infection with *C. difficile* from mice fed either a chow diet (n=9), LF/LF (n=10) or WD (n=14) total in 3 separate experiments (Table S2). Tissue was fixed in 10% formalin in PBS, paraffin embedded and sections cut before hematoxylin and eosin staining by the University of Colorado Histopathology Core. Inflammation was assessed in the cecum using the Barthel scoring system ^24^ and in the colon using the Dieleman scoring system ^23^ by a trained histologist. Briefly, the Barthel system scores damage to the cecum using 0-3 scores for submucosal edema, neutrophil infiltration, number of goblet cells, and epithelial integrity for a composite score of 0 to 12. The Dieleman system scores colonic damage from 0 to 3 for inflammation, and extent of injury, plus scores from 0 to 4 for epithelial regeneration, and crypt damage. Each score is multiplied by a factor from 0 to 4 accounting for % involvement (0 = 0% and 4 = 100%) for a composite score from 0 to 56. Please see original references for more details.

### Statistics

Statistical analyses were performed in R (version 3.4.3 “Kite-Eating Tree”). Data were preprocessed using the “tidyverse” suite ^77^. We used “survminer” and “survival” libraries to analyze mouse survival ^78, 79^. All other data were plotted using “ggplot2”, “ggsignif”, and “cowplot” ^80–82^. All statistical tests were two-tailed with measurements from distinct samples.

### Data availability

The 16S rRNA has been deposited in QIITA ^83^ (Qiita Study ID: 12849) and at EBI (ERP133015).

## Acknowledgments

We would like to thank Jordi Lanis and Sean Colgan for advice on the employed CDI mouse model and Sally Stabler and Whitney Phinney for their assistance in measuring SCFAs. We also appreciate the contribution to this research made by E. Erin Smith, TL(ASCP)CMQIHC, Jenna Van Der Volgen, HT(ASCP)CM, Allison Quador, HTL(ASCP)CM, and Jessica Arnold HTL(ASCP)CM of the University of Colorado for the histology analyses.

## Author contributions

KH conceived of and conducted experiments, analyzed data and co-wrote the paper; CM analyzed data, made figures, and contributed to data interpretation and writing; KA generated and analyzed data from toxin ELISA; NN generated 16S rRNA sequence data and aided in analysis and results interpretation; NMH aided in mouse experiments, 16S rRNA sequencing and generation of toxin ELISA data; NR and MA worked with KH to develop a bile acid panel and aided in analysis and interpretation of results; DO performed histologic evaluation of intestinal tissues; CL directed and contributed to all aspects of the project. All authors contributed to the manuscript.

