## Supplemental Materials for "Dietary fat promotes antibiotic-induced *Clostridioides difficile* mortality in mice"

**Supplementary Materials** for Hazleton K.Z., et al. *Dietary fat promotes antibiotic-induced Clostridioides difficile mortality in mice*

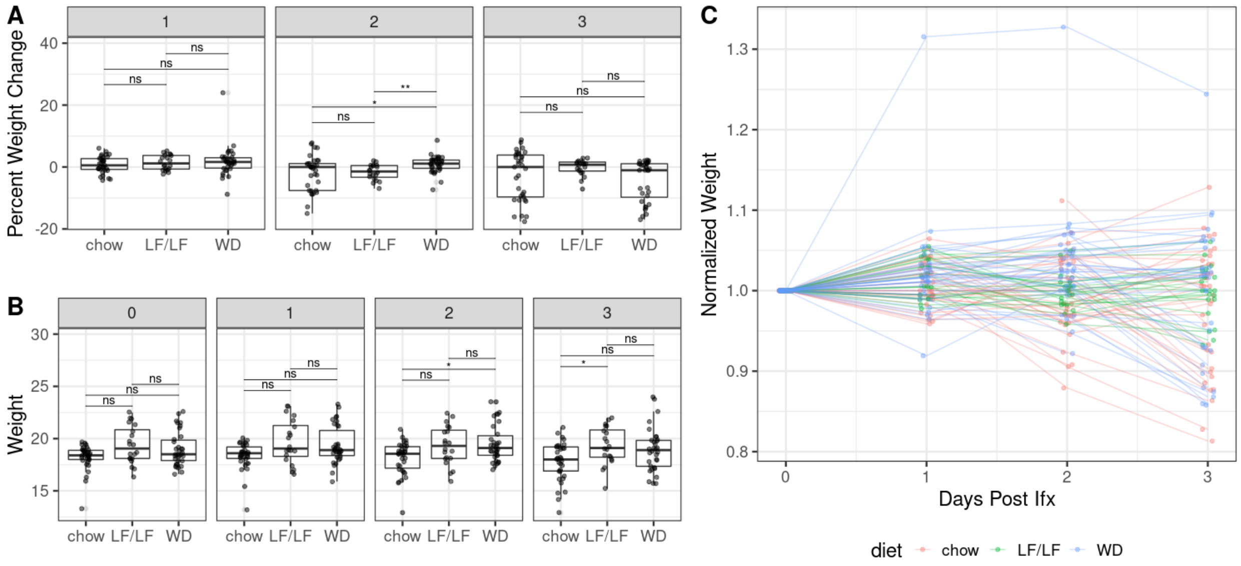


**Figure S1:** Weight of mice before and after infection with *C. difficile* in Cohort 1. (A) Percent weight change of mice after 1,2, and 3 days of infection across diets (baseline was day of infection); (B) weight in grams of mice across diets pre and 1,2, and 3 days post infection and (C) scatter plot of fractional weight change of each mouse, colored by diet. p-values were determined using a Kruskal-Wallis with Dunn's post hoc test. Median and IQR indicated. (* : $p < 0.05$, ** : $p < 0.01$).

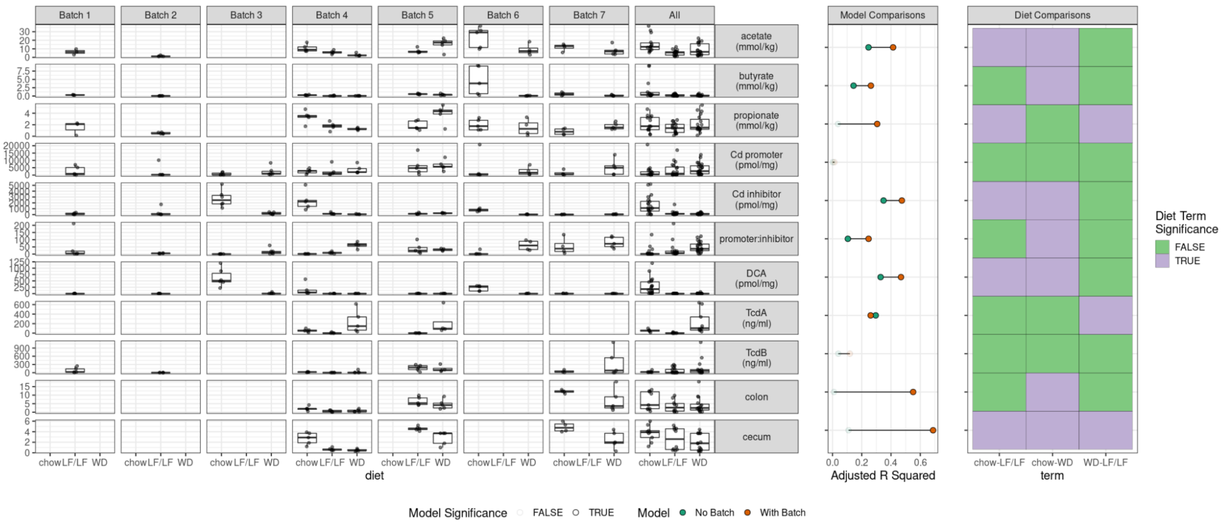


**Figure S2:** Analyte concentrations and inflammatory scores across different batches. The middle panel compares the adjusted R squared of linear models without considering batch (readout ~ diet) to linear models while considering batch (readout ~ diet + batch). Models are opaque if the corrected p values < 0.05 and translucent if not. The right panel shows significant differences between diet groups for models that consider batches.

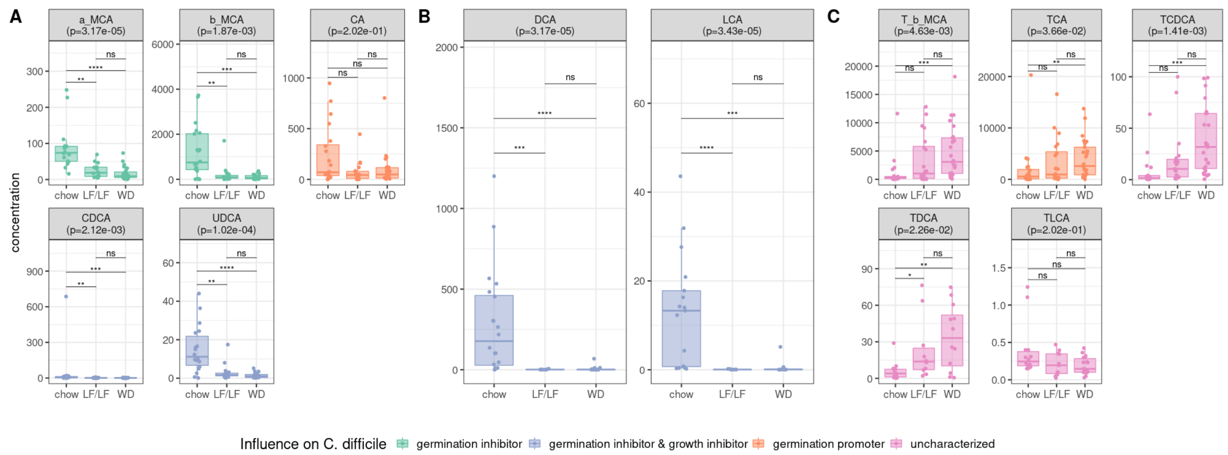


**Figure S3.** Cecal bile acid concentrations (pmol/kg) by diet. Unconjugated primary bile acids (A), secondary bile acid (B) and taurine-conjugated bile acids (C). a_MCA (alpha muricholic acid); b_MCA (beta muricholic acid); CA (cholic acid); CDCA (chenodeoxycholic acid); UDCA (ursodeoxycholic acid); DCA (deoxycholic acid); LCA (lithocholic acid); T_b_MCA (tauro-beta muricholic acid); TCA (taurocholic acid); TCA_3_SO4 (taurocholic acid 3-sulfate); TCDCA (taurochenodeoxycholic acid); TDCA (taurodeoxycholic acid); TLCA (taurolithocholic acid). P-values for the Kruskal-Wallis test with Dunn's post hoc are noted. Median and IQR are indicated. Plots are colored based on the previously described influence of each bile acid on *C. difficile*.

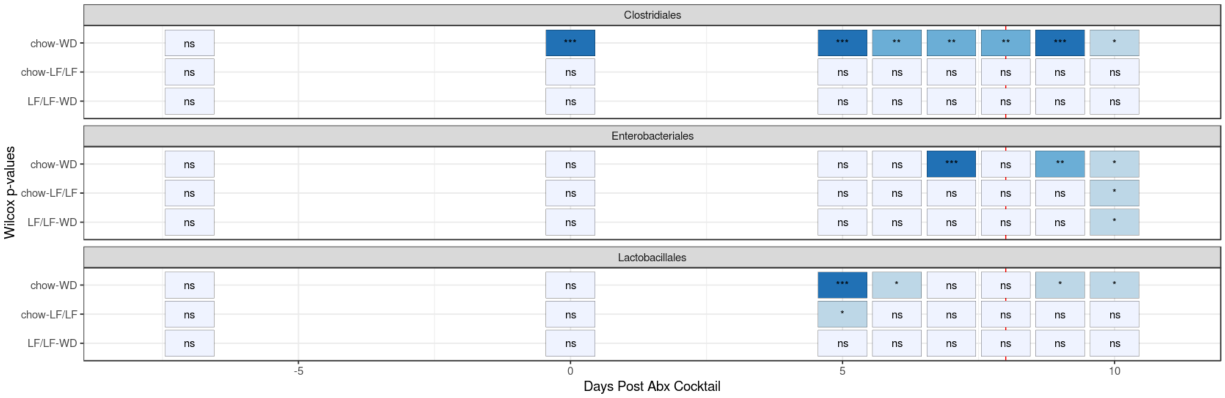


**Figure S4:** Statistical summary of relative abundances of key taxa from Figure 7a. Significant differences between groups are noted as calculated with the Wilcoxon rank-sum test. ***: $p<0.001$. **: $p<0.01$, *: $p<0.05$, ns= non significant.

**Table S1**: Composition of low-fat/low-fiber/high-sucrose diet

| | ***Low-fat/Low-fiber/High-Sucrose*** |
|---|---|
| Fat (% kcal)<br>*(% SFA)*<br>*(% MUFA)*<br>*(% PUFA)* | 17.0<br>*(19.5)*<br>*(41.7)*<br>*(38.8)* |
| Carbohydrates (% kcal)<br>*(Sucrose)* | 64.5<br>*(26.7)* |
| Protein (%kcal) | 18.6 |
| Fiber (g/kg) | 50 (cellulose) |

**Table S2:** Description of samples sizes across different assays in cohorts 1 and 2.

| | | Batches | Chow | LF/LF | WD |
|---|---|---|---|---|---|
| Cohort 1 * (stool collection) | | Total<br>Batch1<br>Batch2<br>Batch3<br>Batch4 | 20<br>10<br>10<br>0<br>0 | 20<br>0<br>10<br>5<br>5 | 20<br>10<br>0<br>5<br>5 |
| | 16S rRNA | Batches 1,2,3 | 13 | 5 | 13 |
| Cohort 2 ** (tissue collection) | | Total<br>Batch1<br>Batch2<br>Batch3<br>Batch4<br>Batch5<br>Batch6<br>Batch7 | 20<br>0<br>0<br>6<br>5<br>0<br>5<br>4 | 20<br>5<br>5<br>0<br>5<br>5<br>0<br>0 | 26<br>0<br>0<br>6<br>5<br>5<br>5<br>5 |
| | histology | Batches 4,5,7 | 9 | 10 | 14 |
| | TcdA | 4,5 | 5 | 10 | 9 |
| | TcdB | 1,2,4,5,7 | 8 | 20 | 15 |
| | Bile acids | All batches | 20 | 20 | 25 |
| | SCFAs | 1,2,4-7 | 14 | 18 | 19 |

*Cohort 1 (longitudinal mortality analysis with serial fecal collection and microbiome sequencing) was conducted in 4 different batches of mice with per batch counts across diets indicated.

**Cohort 2 (tissue collection – intestines for histopathology and cecal aspirates for bile acids, SCFAs, and toxins) was conducted in 7 different batches of mice with per batch counts across diets indicated. 16S rRNA, histology, toxins TcdA and TcdB, bile acids and SCFAs were all measured for particular sets of batches as indicated.

**Table S3**

Complete results of linear modeling statistical analyses performed. **[Table in separate Excel spreadsheet file]**

**Table S4**

| Ingredient (g/KG) | WD | LF/LF |
|---|---|---|
| Anhydrous Milkfat | 36.3 | 50 |
| Beef Tallow | 24.8 | 0 |
| Casein | 190 | 0 |
| Cellulose | 30 | 35 |
| Cholesterol | 0.4 | 0 |
| Choline Bitartrate | 2.1 | 0.01 |
| Corn Oil | 16.5 | 6.5 |
| Corn Starch | 230 | 392.2 |
| L-Cystine | 2.85 | 3 |
| Lard | 28 | 0 |
| Maltodextrin | 70 | 100 |
| Mineral Mix, nTWD (110422) | 35 | 15 |
| Olive Oil | 28 | 16.5 |
| Sodium Chloride | 4 | 0 |
| Soybean Oil | 31.4 | 23 |
| Sucrose | 255.6 | 24 |
| TBHQ, antioxidant | 0.028 | 0.014 |
| Thiamin (81%) | 0.015 | 0.002 |
| Vitamin K1, phylloquinone | 0.003 | 0.014 |
| Vitamin Mix* | 15 | 2.75 |

* WD used the Vitamin Mix nTWD (110423) and LF/LF diet used AIN-93-VX (94047)
